# Juvenile niches select between two distinct development trajectories and symbiosis modes in vent shrimps

**DOI:** 10.1101/2023.07.02.547428

**Authors:** Pierre Methou, Marion Guéganton, Jonathan T. Copley, Hiromi Kayama Watanabe, Florence Pradillon, Marie-Anne Cambon-Bonavita, Chong Chen

## Abstract

Most animal species have a singular developmental pathway and adult ecology, but developmental plasticity is well-known in some like honeybees where castes display profoundly different morphology and ecology. An intriguing case is the Atlantic deep-sea hydrothermal vent shrimp *Rimicaris hybisae/chacei* that share dominant COI haplotypes but develops into either the symbiont-reliant *hybisae* with a hypertrophied head chamber (in the Mid-Cayman Spreading Centre) or the mixotrophic *chacei* with a narrow head chamber (on the Mid-Atlantic Ridge). Here, we use X-ray micro-computed tomography and fluorescent *in situ* hybridization to show that key anatomical shifts in both occur between the juvenile-subadult transition, when those developing into *hybisae* have fully established symbiosis but not *chacei*. On the Mid-Atlantic Ridge the diet of *R. chacei* has been hypothetically linked to competition with the obligatorily symbiotic congener *R. exoculata*, and we find anatomical evidences that *R. exoculata* is indeed better adapted for symbiosis – suggesting the *chacei* morph could be an adaptation to prevent competitive exclusion. Our results suggest that the two distinct development trajectories are likely selected according to the diet available to the juveniles, determined by whether energetically sufficient symbiont colonisation has occurred before reaching subadult stage.

## Introduction

Each animal species is generally constrained to a single mode of ontogenetic development, leading to individuals (of the same sex) exhibiting similar ecology at each life stage. Nevertheless, there are some well-documented cases where individuals of the same species can develop into profoundly different forms with distinct ‘ways of life.’ Castes of eusocial insects is a well-known example, where nutritional input typically has a key role in determining the development outcome. Whether a female larva of the honeybee *Apis mellifera* develops into worker or queen, for example, is almost entirely dependent on if it is fed a large quantity of royal jelly [1]. Another example is the cave fish *Astyanax mexicanus*, whose morphology differs greatly among individuals living in rivers and surrounding caves – cave morphotypes have lost their eyes and show reduced pigmentation but have larger body sizes and other sensory organs [2]. At hydrothermal vents, a significant case is the *Ridgeia piscesae* tubeworm, which develops into a short-fat or a long-skinny morphotype depending on the vent flow regime [3,4].

The formation of symbioses, intimate and long-term biological association between two or more organisms [5], is also a powerful evolutionary force that can have fundamental impacts on the developmental processes and ecology of the host animal [6–8]. For example, the bobtail squid *Euprymna scolopes* houses the bioluminescent symbiont *Vibrio fisheri* for counter-illumination in its light organ which morphogenesis requires signalling molecules from the symbiont [9]. Deep-sea hydrothermal vent ecosystems are unusual in that intimate symbiotic relationships between chemoautotrophic bacteria and endemic invertebrate animals form the basis for the entire system [10]. Vent holobionts usually acquire their symbionts from the environment after they settle in the vent field as juveniles – with the exception of vesicomyid clams which have maternal inheritance of chemosynthetic symbionts [11]. As significant transitions in ecological niche within the life cycle of a species often involve metamorphosis, where conspicuous anatomical and physiological shifts occur, symbiont acquisition is often linked to such metamorphosis involving the loss or the reduction of the digestive organs and an enlargement of the symbiont hosting structures [12–14]. These reconfigurations of internal organs post-settlement are not necessarily apparent from the external morphology, as seen with the ‘cryptometamorphosis’ of *Gigantopelta* vent snails where the symbiont-hosting ‘trophosome’ suddenly expands internally [15] or the straightening of the digestive tube in bathymodioline mussels [16].

Among vent animals, alvinocaridid shrimps exhibit a wide range of feeding strategies and reliance on symbiosis. A strict dependence on symbiosis is found in two closely related species, *Rimicaris exoculata* and *R. kairei*, hosting dense communities of filamentous bacterial symbionts inside their expanded ‘head’ (the cephalothorax) that provides most of their diet [17–20]. These symbiotic species exhibit striking morphological differences from other alvinocaridid shrimps, with the enlargement of their symbiont-hosting cephalothoracic cavities and mouthparts starting during the post-settlement metamorphosis between juveniles and adults [19,21]. Conversely, several other species of vent shrimps such as *Nautilocaris saintlaurentae* and *Rimicaris variabilis* harbour few to no symbiotic bacteria and instead feed on other food sources such as bacterial mats or detritus [22,23].

One species complex of alvinocaridid shrimps from Atlantic vents – the *Rimicaris hybisae/chacei* complex – is exceptional as *R. hybisae* and *R. chacei* share dominant *COI* haplotypes (electronic supplementary material, figure S1) but metamorphose into two distinct forms occupying disparate ecological niches depending on the environment [24,25]. The apparent differences in morphology led to their descriptions as separate species [26], but currently available genetic barcode data do not support the existence of two distinct species-level clades [24,27]. *Rimicaris hybisae* occurs in the Mid-Cayman Spreading Centre (MCSC), mostly rely on symbiosis and has an external morphology resembling *R. exoculata* with an inflated cephalothorax [26,28]. It appears to still retain an ability to feed on alternative sources outside of its symbiosis, with evidences of facultative carnivory/scavenging in individuals distributed at the periphery of vent fluid emissions [25]. In contrast, *R. chacei* from the Mid-Atlantic Ridge (MAR) co-occur with *R. exoculata* [29] and lacks an enlarged cephalothorax; it has a mixed diet where it also feeds on bacterial mat and scavenges in addition to some contribution from symbiosis [21,30].

Although *R. exoculata* and *R. chacei* co-occur at MAR vents, *R. exoculata* dominates the vigorously venting ‘black-smoker’ chimneys whereas *R. chacei* mainly occurs in diffuse flow environment with much weaker fluid input [29]. This is in stark contrast to the situation at MCSC where *R. hybisae* is the only shrimp species dominating the surface of active smoker chimneys [26]. This leads to a hypothesis that the *R. chacei* adult morphology is a result of competitive exclusion during the juvenile stage from the vigorously venting environment by *R. exoculata* [29], and that this environment is necessary for development into the *R. hybisae* morphology. Anatomy and food source are closely linked in animals, and organ volumetric is a useful estimator of feeding ecology for hydrothermal vent species where detailed *in situ* studies are difficult [31]. Until now, no direct comparisons in the anatomy, ontogenetic changes, and symbiont acquisition between *R. hybisae* vs *R. chacei* have been made, or how they compare to the obligatorily symbiotic *R. exoculata*. In order to shed light on this hypothesis, we combine quantitative 3D anatomy by X-ray micro-computed tomography (µ-CT) and fluorescence in situ hybridisation (FISH) to answer three questions: 1) At which life stage does the split in development trajectories occur in *R. hybisae/chacei*, 2) Whether this is correlated with symbiont acquisition timing, and 3) Is there anatomical evidence that *R. hybisae/chacei* would indeed be outcompeted by *R. exoculata*.

## Methods

Specimens of the three main target shrimp species – *R. exoculata, R. hybisae*, and *R. chacei* – were collected at hydrothermal vent fields during several sea-going expeditions: *R. exoculata* and *R. chacei* were collected at the TAG and Snake Pit vent fields during the BICOSE 2 (R/V *Pourquoi pas?*, 2018, doi: https://doi.org/10.17600/18000004) expedition using suction sampler of the human-occupied vehicle (HOV) *Nautile*; *R. hybisae* were collected at the Von Damm and Beebe vent fields during the expeditions JC82 (RRS *James Cook*, 2013) and YK13-05 (R/V *Yokosuka*, 2013) expeditions, respectively, with suction samplers on ROV *Isis* or HOV *Shinkai 6500*. In addition, we also studied the anatomy of *R. kairei* to confirm that it does not differ significantly from that of its closest relative, *R. exoculata*; using specimens collected from the Kairei and Edmond vent fields during expeditions YK09-13 (R/V *Yokosuka*, 2009) and YK16-E02 (R/V *Yokosuka*, 2016) using suction samplers mounted on the HOV *Shinkai 6500*. Shrimps were preserved in 10% formalin or 99% ethanol and measured with vernier calliper using carapace length (CL) for standardized measurement, upon recovery on-board. Individuals were identified to their life stages, according to Methou *et al*. [21]. Subadults of all species were characterized as small individuals morphologically similar to adults with sizes below the onset of sexual maturity whereas stage A and stage B juveniles of *R. exoculata* were distinguished by the morphology of their eyes [21]. Given that *R. chacei* juveniles settle and begin their anatomical transitions at smaller sizes than *R. exoculata*, their adult stages were separated into two categories: ‘small adults’ of similar sizes to that of *R. exoculata* juvenile stages (CL = 8-10 mm) and ‘adults’ (CL= 14-16 mm). Life stages of *R. kairei* was defined following the established nomenclature for *R. exoculata* and the same was done for *R. hybisae* based on *R. chacei*. Additional subadult specimens of *R. exoculata, R. chacei*, and *R. hybisae* were fixed in 3% paraformaldehyde (PFA) for 3 hours on board, rinsed with phosphate buffered saline (PBS) / sterile seawater buffer and stored in 50:50 2x PBS/ethanol at −20^⁰^C for fluorescence *in situ* hybridization (FISH) analyses (electronic supplementary material, table S1).

A total of 47 individuals, including a triplicate of each life stages for the four-shrimp species (except where only two individuals were available for stage A juveniles of *R. hybisae*), were used for X-ray micro-computed tomography (µ-CT) scanning. Shrimps preserved in 99% ethanol were progressively rehydrated in Milli-Q water in four steps (25:75, 50:50 and 75:50 Mili-Q: Ethanol), and then stained for 24 hours in 0.05 mol/L iodine solution. Scanning was performed using a ScanXmate-D160TSS105 (Comscantecno, Japan) commercial μ-CT at 60-90 kV/25-92 µA, with a resolution of 10.904-49.587 µm per pixel, at 992 × 992 pixels resolution per slice (for details see electronic supplementary material, table S2). Images obtained were imported into the specialist software Amira version 2022 (Thermo Fisher) for visualisation and manual segmentation of the different organs of interest. This included the symbiont-hosting branchiostegite (i.e., wall of the cephalothoracic gill chamber) and mouthparts (scaphognathites and exopodites), as well as the digestive system (stomach, digestive tube, and hepatopancreas). Only the anterior part of the cephalothoracic cavities were reconstructed, since posterior components facing the gills are known to be not colonized by bacterial symbionts [18,20,30]. Prior to smoothing, organ volumes and surface areas were calculated also in Amira based on the segmentation. As pereopods and antennae were often fully or partially broken, these parts were removed from all our 3D reconstructions and thus the measurement of total body volumes and surface areas (see Table S3 for detailed body volumes and surface areas measurements and supplementary material for 3D interactive models). The final surface renderings used for the figures were generated post-smoothing and complexity-reduction. Cephalothoracic cavities of five specimens at each life stage of *R. chacei* and *R. hybisae* (except where only two individuals available for stage A juveniles of *R. hybisae*) were also dissected to observe the distribution of bacteriophore setae on their mouthparts. Mouthparts of one *R. exoculata* adult were also dissected and observed. As the anatomical features and patterns seen in *R. kairei* generally matched *R. exoculata*, for simplicity and clarity we only displayed the three main study species in the main figures and the *R. kairei* data are figured in electronic supplementary material, figures S2-S4.

For FISH analyses of subadult symbiont colonization, the cephalothorax of *R. exoculata* and *R. chacei* subadults were progressively dehydrated in a PBS-ethanol series (ambient temperature) before embedding in polyethylene glycol distearate-1-hexadecanol (9: 1) resin (Sigma, St. Louis, MO). Resin blocks were stored in −20°C until trimming into 10 µm sections with a RM 2255 microtome (Leica Biosystems, Nussloch, Germany). Sections were placed on adhesive glass slides (*Menzel-Gläser Superfrost® Plus, USA)*, residues of resin were removed prior to hybridization with three baths of 96% ethanol for 5 min each and *tissues were partially rehydrated with a bath of 70 ethanol*. The cephalothorax of *R. hybisae* subadults were cut from the abdomens and rehydrated in 50:50 PBS-ethanol bath followed by 75:25 PBS-ethanol (30 min each) and then rinsed three times in 1x PBS buffer for 5 min each. Specimens were then progressively transferred to an optimum temperature compound (OCT) (Tissue-Tek; Sakura Finetek, Osaka, Japan): first in 15% sucrose-PBS, followed by 30% sucrose-PBS, and finally in 30% sucrose-PBS/OCT, for 1h 30 min each, before embedding in OCT in a plastic holder at 4°C. Sectioning of OCT-embedded specimens was done with a cryostat (CM1520; Leica Biosystems) in a chamber at −20°C. Sections of 8 µm thickness were thaw-mounted on adhesive glass slides. Before hybridization, mounted sections were washed in 1x PBS two times for 5 min each, then postfixed 1x PBS-10% formalin for 10 min, and finally washed again in 1x PBS for three times at 5 min each. These two different embedding and sectioning protocols have already been used for FISH experiments with *Rimicaris* shrimps and are not expected to impact the detection of symbionts [18,19].

For all species, sections were hybridized in a mix containing 0.5 mM (0.5 μM final concentration) of the Eub338 (eubacteria) probe in a 30% (*R. chacei* & *R. exoculata*) or 35% (*R. hybisae*) formamide hybridization buffer for 3 h at 46°C following a published protocol [19]. Sections were washed at 48°C for 30 min in a washing buffer [19], rinsed briefly with miliQ water, air dried, and covered with SlowFade Gold antifade reagent mounting medium containing DAPI (Invitrogen) and a cover slip. Observations of *R. exoculata* and *R. chacei* subadults were made using a Zeiss Imager.Z2 microscope equipped with the Apotome.2® sliding module and Colibri.7 light technology (Zeiss, Oberkochen, Germany) and processed with the Zen software (Zeiss). Observations of *R. hybisae* subadults were made using a Leica Stellaris STED confocal scanning microscope (Leica, Germany) and analysed with the LAS X software.

## Results

Anatomy of *Rimicaris* species was reconstructed across four stages of development from recently settled juvenile to adult stages (figure 1; electronic supplementary material, figures S2 & S3). Early juvenile stages of all species had very small stomach volumes, ranging from 0.12 ± 0.01% of the total body volume in *R. exoculata* to 0.32 ± 0.04% in *R. chacei* (Figures 1 and 2). In adults, *R. chacei* and *R. hybisae* both possessed large stomach volumes representing 1.73 ± 0.84% and 2.18 ± 1.3% of their body volumes, respectively (Figures 2 and 3); whereas adult stages of *R. exoculata* had much smaller stomachs at 0.69 ± 0.32% of the body volume (Figures 2 and 3). However, their stomachs expanded differently during their transition from juvenile to subadult with a dramatic increase, although variable among individuals, up to 2.03 ± 0.99% right from the subadult stages of *R. chacei* while it was more gradual in *R. hybisae* with an increase up to 0.86 ± 0.09% in subadults and 0.77 ± 0.13% in small adults (Figure 1 and 2). On the other hand, the hepatopancreas followed a similar trajectory in the three species, occupying a very large volume in juvenile stages (*R. exoculata*: 18.6 ± 1.3%; *R. hybisae*: 17.2 ± 0.96%; *R. chacei*: 17.3 ± 4.0%) representing lipid reserves, and then decreasing rapidly when metamorphosising into subadults (*R. exoculata*: 8.55 ± 1.93%; *R. hybisae*: 6.61 ± 1.07%; *R. chacei*: 8.02 ± 2.85%). No clear variation could be seen in the volumes of digestive tubes along the metamorphosis of the three species (figures 2 and S4). Overall, the anatomical development of the *R. kairei* digestive system exhibited similar patterns as seen in *R. exoculata*, although with an earlier stomach enlargement in stage B juveniles (*R. kairei*: 0.60 ± 0.26%; *R. exoculata*: 0.15 ± 0.04%) as well as an earlier decrease of hepatopancreas volume (Juveniles B; *R. kairei*: 8.08 ± 3.23%; *R. exoculata*: 18.6 ± 1.3%) (electronic supplementary material, figures S3-S4).

**Figure 1.**
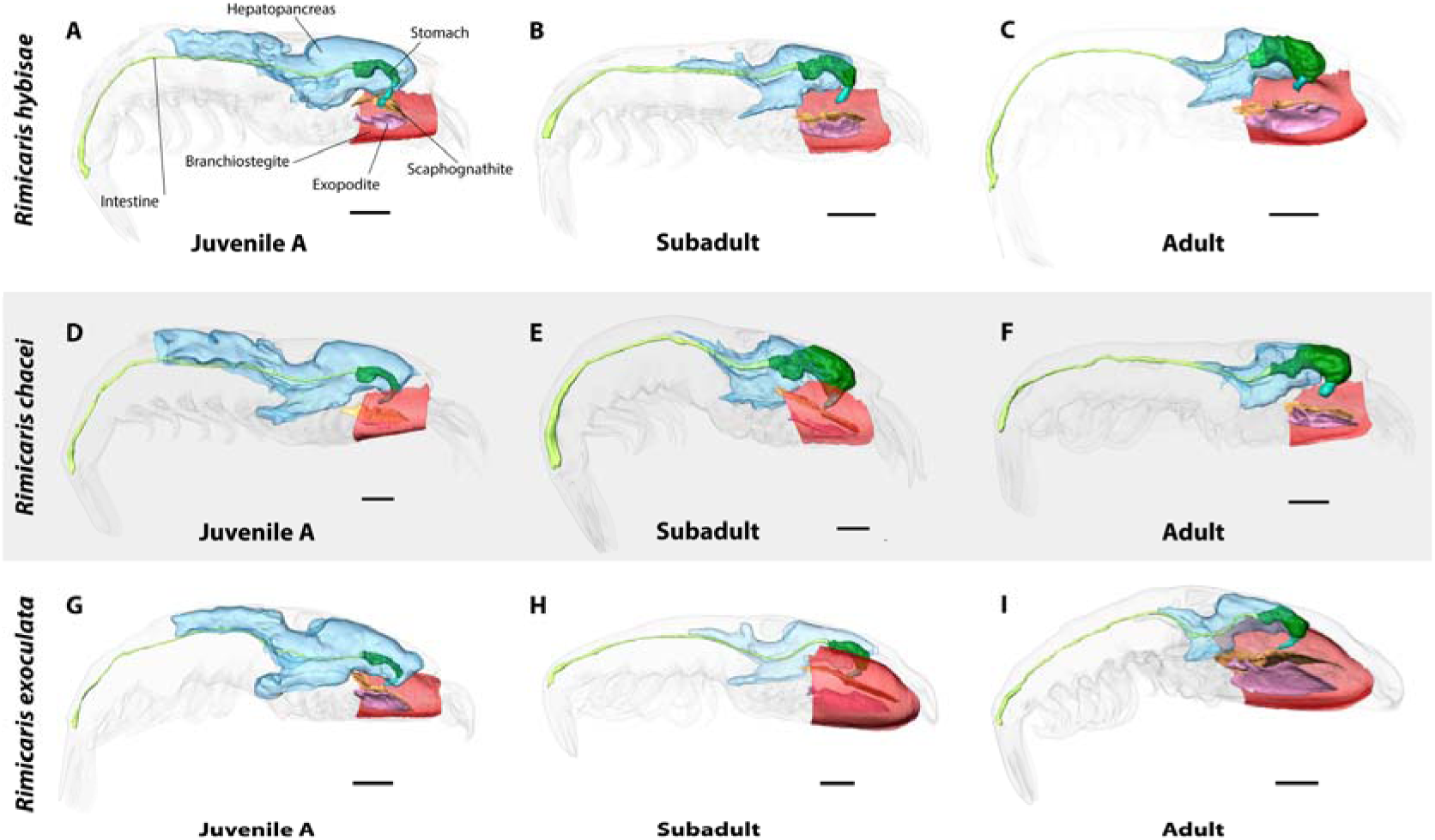
3D anatomical reconstructions of symbiont hosting organs (branchiostegite, scaphognathites, and exopodites) and digestive organs (stomach, hepatopancreas, digestive tube) in three *Rimicaris* shrimps across post-settlement ontogeny of (A-C) *R. hybisae*, (D-F) *R. chacei*, (G-I) *R. exoculata*.

**Figure 2.**
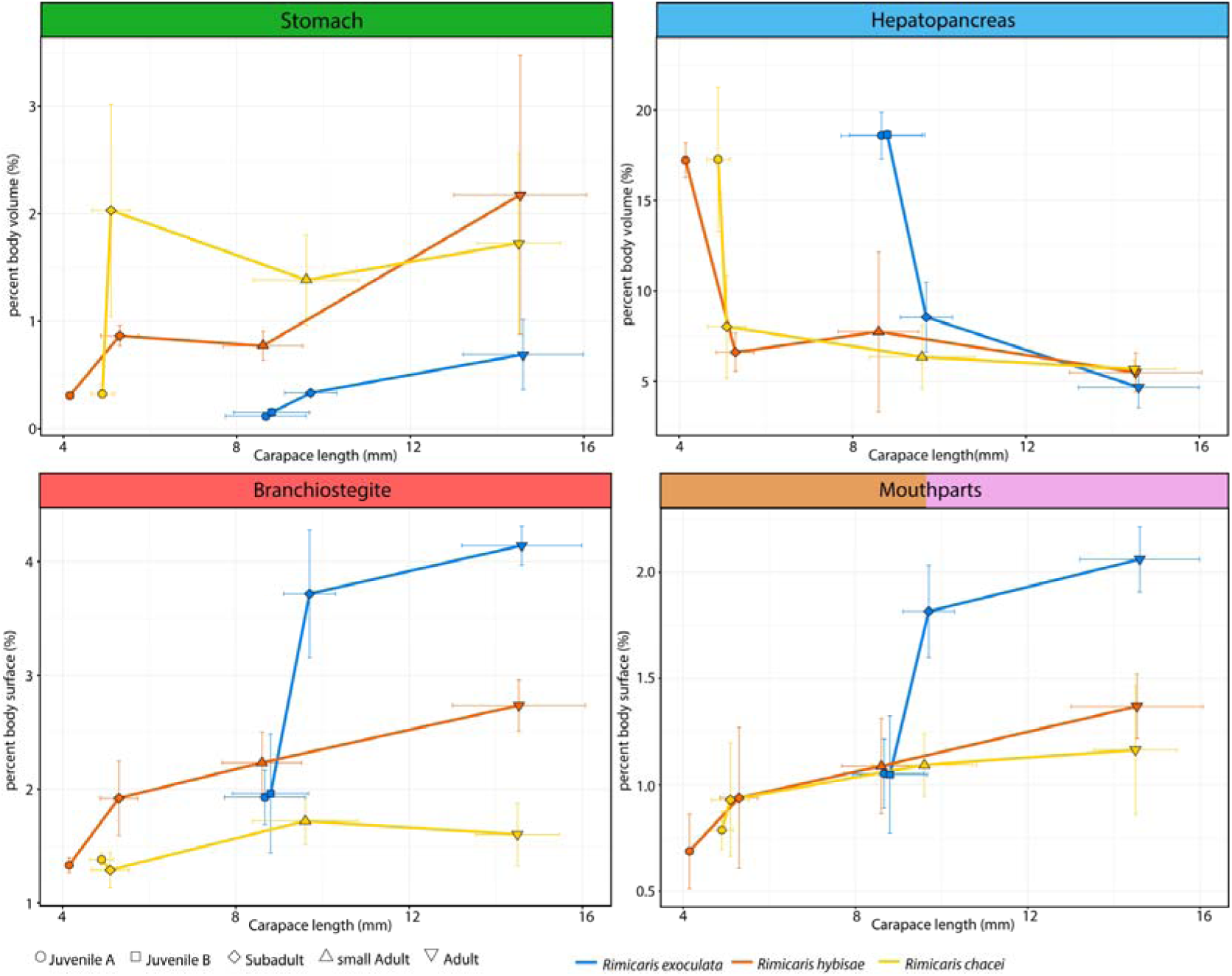
Relationship between body size (carapace length) and the relative per cent body volumes or body surfaces of symbiont-hosting organs (branchiostegite, mouthparts) and digestive organs (stomach, hepatopancreas).

**Figure 3.**
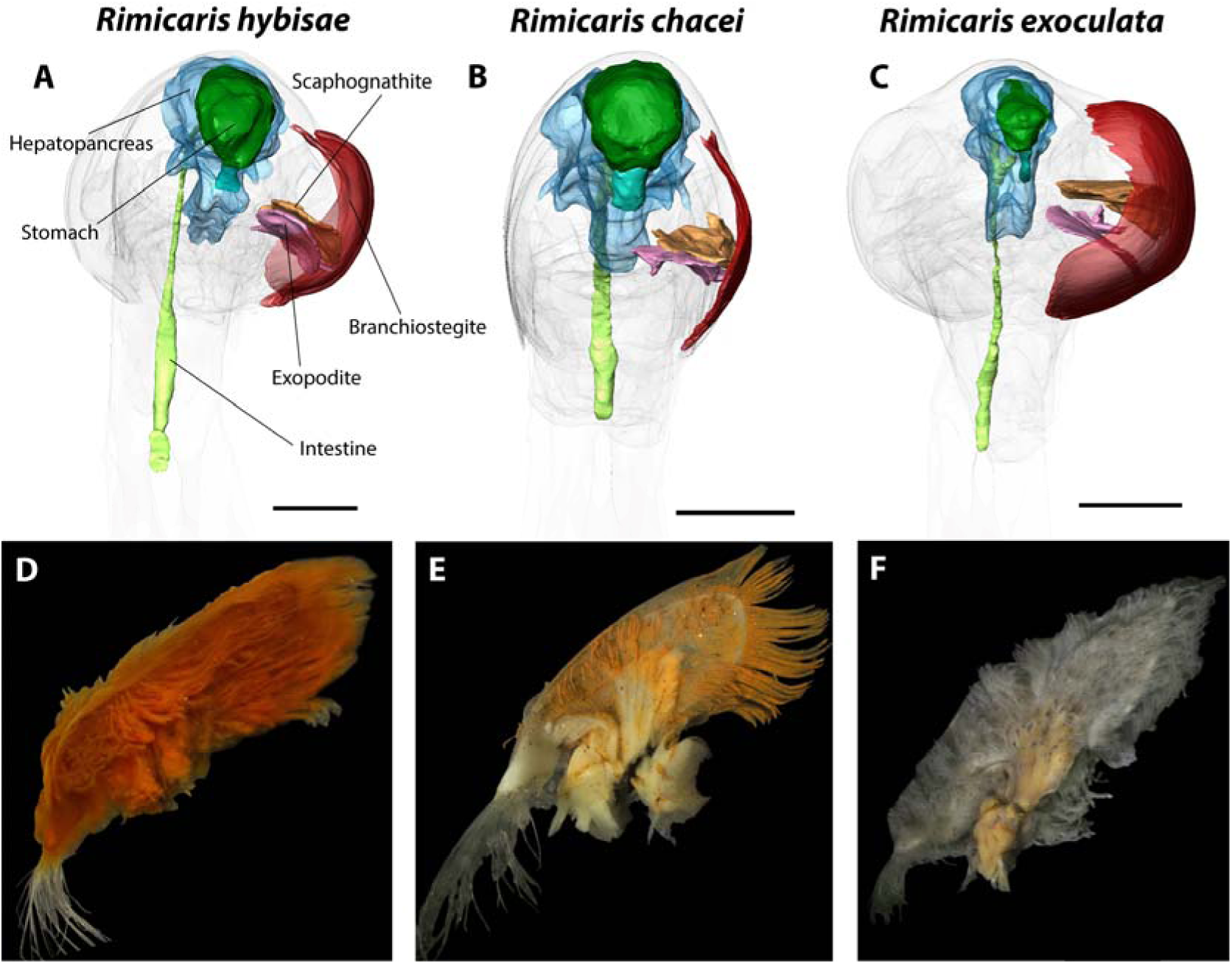
Symbiont hosting organs in adult stages of the three *Rimicaris* shrimps. (A-C). 3D reconstructions of hosting organs in frontal view. (D-F) Dissected mouthparts covered with bacteriophore setae.

The symbiont-hosting branchiostegite (wall of the gill chamber) surface area was found to exhibit a sudden expansion in *R. exoculata* between juvenile B (1.96 ± 0.52% of total body surface) and subadult stages (3.72 ± 0.56% of the total body surface), and then up to 4.14 ± 0.17% in adult stages (figures 1 and 2). Surfaces of other symbiont-hosting mouthparts – scaphognathites and exopodites – similarly expanded between juvenile B and subadult stages of *R. exoculata* from 1.05 ± 0.28% to 1.82 ± 0.22% of the total body surfaces (Figure 2). The pattern was also seen in *R. kairei* with rapid expansion before subadult, main differences being in *R. kairei* the expansion was also evident between juveniles A and B and the expansion being less and more variable in adults (see electronic supplementary material, figures S3-S4). The increase of branchiostegite surfaces was more rapid in *R. hybisae* between juvenile and subadult stages and more or less isometric thereafter, from 1.33 ± 0.06% of the total body surface in stage A juveniles to 1.92 ± 0.33% in subadults and 2.74 ± 0.23% in adults. Conversely, surfaces of *R. chacei* branchiostegites expanded only slightly along their ontogeny from 1.38 ± 0.05% to 1.60 ± 0.05%. For mouthparts, the increment was less marked than *R. exoculata* for both *R. hybisae* and *R. chacei*, with a limited growth of both mouthparts from 0.69 ± 0.18% to 1.37 ± 0.15% between stage A juveniles and adults in *R. hybisae* and 0.79 ± 0.09% to 1.16 ± 0.30% in *R. chacei* (Figure S4). Dissection of the adult mouthparts between *R. hybisae* and *R. chacei* however, showed a striking difference in the coverage of bacteria-hosting setae; being much less dense on *R. chacei* mouthparts than on *R. hybisae* and *R. exoculata* mouthparts (figure 3). The dense setae would mean a much greater colonisable surface area in *R. hybisae* mouthparts than *R. chacei* (figure 3). Our FISH observations of subadult specimens indicate that the three taxa differed in the timing of their complete bacterial colonization (figure 4). Although subadult mouthparts were all colonized by filamentous bacteria in the three species, the branchiostegites of *R. exoculata* and *R. hybisae* subadults were densely colonized across the entire surface, but only partly in *R. chacei*.

**Figure 4.**
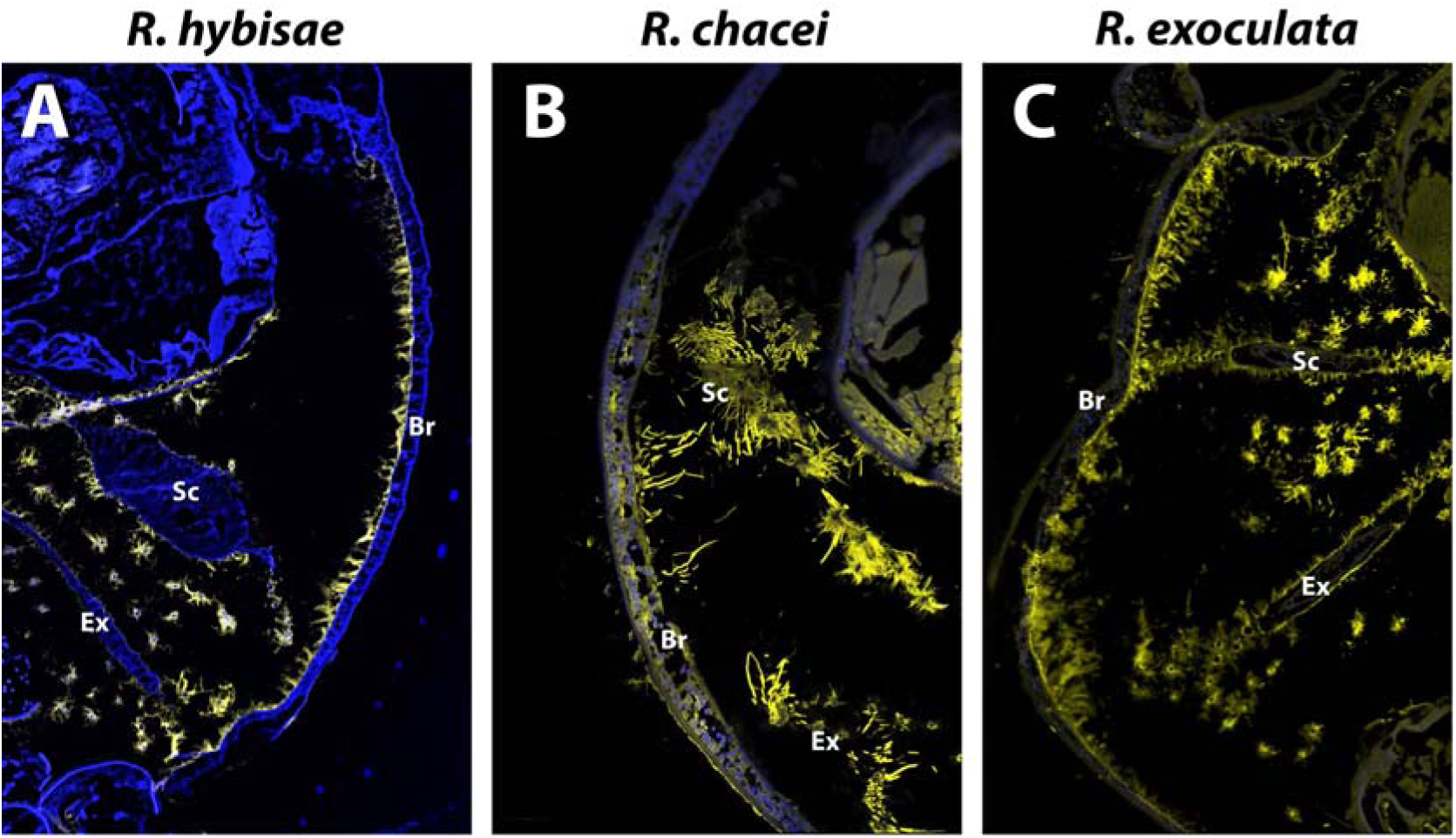
FISH observations of cephalothoracic cavities of *Rimicaris* subadults with universal bacterial probes (yellow) on semi-thin sections (7µm) stained with DAPI (blue). A. *R. hybisae* subadult B. *R. chacei* subadult C. *R. exoculata* subadult

## Discussion

Our results confirm that *Rimicaris hybisae* and *R. chacei* indeed display striking differences in anatomical development, despite the lack of genetic differentiation based on mitochondrial *COI* gene suggesting they are likely two morphotypes of the same species-level lineage [24,27]. The symbiont-hosting organs of the *R. hybisae* form increased in percentage surface area (as a fraction of the overall body surface) throughout growth with the fastest increase between juvenile and subadult stages, whereas there was almost no increase in *R. chacei*. The symbiont-hosting mouthparts were not different in the overall surface area between the two forms, but they were densely covered by setae only in *R. hybisae* (like *R. exoculata*) meaning a much higher realised surface area for colonisation in *R. hybisae*. The *chacei* morphotype also exhibits a dramatic increase in stomach size before reaching subadult stage, whereas this was not seen in *R. hybisae*. As such, the most significant shifts in the volume or surface area of key organs (figure 2) was clearly between juvenile and subadult stages, indicating that the development trajectory into either form is determined in the ‘critical period’ between these two stages. All species examined rapidly decreased in the volume of hepatopancreas before reaching the subadult stage, reflecting a rapid depletion of the lipid reserve accumulated before settlement [21]. This means the shrimps must become energetically self-sustaining before reaching the subadult stage. Taken together, these results strongly imply that the ecological niche occupied by juvenile *R. hybisae/chacei* before reaching subadult size is key in deciding the development trajectory and therefore the niche occupied as adults. The total anatomical shift between juvenile and subadult stages may be sufficient to be considered a post-settlement metamorphosis [15].

Still, there are two key anatomical characters shared across *R. hybisae* and *R. chacei* that are distinct from the anatomical characteristics of *R. exoculata* (and *R. kairei*). The final stomach volumes of *R. hybisae* and *R. chacei* are equivalent at the adult stage, both being much larger than that of *R. exoculata*. The overall mouthpart surface areas are also smaller in both *R. hybisae* and *R. chacei* compared to *R. exoculata*. Furthermore, large variations in stomach volumes of adults could be seen among individuals of both *R. hybisae* and *R. chacei*. This inter-individual variability could be related to plasticity in feeding ecology with the potential for facultative carnivory/scavenging reported for some *R. hybisae* individuals [25] or mixed diets with variable contribution from food and symbiosis among *R. chacei* individuals [21,30]. We recognise that potentially different fixation conditions of specimens collected during different expeditions could be responsible for the observed variability to some extent, but this should not influence the general trends observed. Thus, although *R. hybisae* is much more similar to *R. exoculata* superficially at a glance from the external morphology, some features of its internal anatomy are much more similar to *R. chacei* and are indicative of their genetic relationship (electronic supplementary material, figure S1) [24,27].

Enlargement of symbiont-hosting organs and the correlated reduction of digestive system are prime anatomical signals in hot vent species for increased reliance on chemosymbiotic bacteria, resulting in increased energetic yield [31,32]. In the cephalothorax, the surface areas of all symbiont-hosting organs in *R. exoculata* exhibited sudden, exponential growth between stage B juveniles and subadults like between juvenile A and subadults for *R. hybisae*, but to a much greater extent. The larger symbiont-hosting organs coupled with reduced stomach volume in *R. exoculata* compared to *R. hybisae/chacei* are therefore anatomical evidences that *R. exoculata* is indeed the more specialised species for chemosynthetic symbiosis and would therefore outcompete the less specialised *R. hybisae/chacei*, as previously hypothesised [17,18,25,28]. Since *R. exoculata* co-occurs with *R. chacei* on MAR vent fields, juveniles of the two species have been suggested to directly compete for space and thus access to the vent fluid source, with *R. exoculata* outcompeting *R. chacei* from the hottest and most energetically superior habitable zone on active smoker chimneys [29]. This is supported by their spatial segregation patterns in cooler areas for *R. chacei*, and clear differences in their population demographics with a collapse of *R. chacei* populations following settlement phase suggestive of a high juvenile mortality [29]. This interspecific competition with *R. exoculata* may limit acquisition of symbionts for *R. chacei* juveniles situated away from the influence of vent fluid and ultimately hamper the development of an energetically sufficient chemosymbiosis; promoting a shift towards a mixotrophic feeding strategy [29]. On the other hand, *R. hybisae* has no known competitor at the MCSC vents [24,33] and can fuel its chemosymbiosis from its early acquisition in juvenile stages. This niche differentiation between juveniles of *R. hybisae/chacei* in MCSC vs MAR could therefore drive the separation of local individuals onto two ontogenetic development pathways.

Our observations show that *R. hybisae* and *R. chacei* differ in the timing of their symbiont colonization on branchiostegites, being densely colonized in the subadult stages in *R. hybisae* but not *R. chacei*, despite the organ being colonised by bacteria in the adults of both forms [26,30]. This coincides well with the timing of key anatomical shifts before the subadult stage, and suggests having well-colonized branchiostegites during this ‘critical period’ may be a condition for taking the symbiosis-reliant developmental trajectory. The delayed symbiont colonisation in *R. hybisae/chacei* at MAR vents during this competency window, due to being outcompeted by *R. exoculata* and potentially unable to provide sufficient chemosynthetic energy sources for its symbionts, may lead to individuals going down the mixotrophic *R. chacei* pathway [21,29]. As such, symbionts may drive the morphogenesis of enlarged symbiont-hosting organ in the shrimp. Though this is still speculative in our case, similar cases are known, such as the morphogenesis of the light organ in juveniles of the bobtail squid *Euprymna scolopes* which are mediated by the recognition of microorganism-associated molecular patterns (MAMPs) from its *Vibrio fischeri* symbionts upon acquisition [9]. Moreover, *E. scolopes* juveniles treated with antibiotics enter in a refractory state after five days and cannot be recolonized by *V. fischeri* symbionts [34]. In juvenile *Rimicaris* shrimps, the transition from juvenile to subadult is certainly aided by red lipid reserves stored in the hepatopancreas and the depletion of this may mark the end of the ‘critical period.’ This would suggest that shifts in the cephalothorax anatomy of *R. chacei* and *R. hybisae* are reversible either to a more or a less developed symbiosis if exposed to different conditions during their juvenile development.

In species with differential adult phenotypes and therefore developmental pathways, an individual’s fate could be determined purely by environmental factors or genetic factors, or anything in the spectrum in-between [1]. For example, phenotype variations in blind cavefishes appear to be purely genetic [2,35] whereas caste determination in different species of eusocial insects stem from a combination of various environmental stimuli and genetic basis mediated by epigenetics [1]. The two distinct morphotypes of *Ridgeia piscesae* tubeworms is a similar case in vents, influenced by vent flow dynamics [3]. This rapid divergence among populations can be facilitated by phenotypic plasticity or by standing genetic variation with pre-existing polymorphism of relevant alleles in ancestral populations [36–38]. Freshwater populations of the stickleback fishes *Gasterosteus aculeatus* which recently and repeatedly diverged from marine ancestors in several rivers and lake around the world after the last glaciation [39] exhibit a high level of phenotypic variability with the freshwater ones exhibiting reduction in bony plates linked to an allele that was already present at low frequency in the ancestral population [37,39]. On the other hand, divergences among streams and lakes ecotypes of this same species in terms of body depth, head shape, or morphology of the digestive system, were found to be related to an interplay of phenotypic plasticity and genetic predisposition [40]. Though we hypothesise that the developmental fate of *R. hybisae/chacei* is largely environmental and down to accessibility to the active smoker habitat and the timely colonisation of symbionts on the branchiostegites, fine-scale genomic data contrasting the two forms is still missing. Therefore, we cannot exclude the possibility of genetic factors having important roles as well. Future work, including population genomics, will be required to disentangle the respective influence of environment vs genetics on the phenotype and ecology of these shrimps.

Understanding how chemosymbiotic relationships are established is a major challenge because symbiosis is such a successful and effective strategy that most hosts energetically reliant on symbionts have become obligate holobionts that cannot survive or develop into adulthood to reproduce without their partners [13,41]. Our results highlight *R. hybisae/chacei* as an exceptional case where the host is able to take a separate, mixotrophic development trajectory if symbiont colonization is limited, as a ‘back up plan’ to survive and reproduce. We show that individuals have a limited ‘critical period’ before reaching subadult stage that determines the trajectory, likely by whether an energetically viable symbiosis has been established by then. Importantly, with this new information, *R. hybisae/chacei* can be a valuable model for understanding how chemosymbioses are established and how they function, as it can provide comparable data for genetically equivalent individuals in different symbiotic conditions and also during transitions – like how the blind cavefish model has proved useful for studying eye loss [35]. Our work therefore lays the foundations for studying how symbiotic relationships emerge and how dialogues can be established between animals and microorganisms.

## Supporting information

supplemental_3Dmodels

## Acknowledgments

We thank the captains and crews of the different research cruises conducted on-board R/V *Pourquoi pas?* (BICOSE 2), R/V *Yokosuka* (YK09-13, YK13-05 & YK16-E02), and R/V *James Cook* (JC82). We also thank the pilots and the operation team of the HOV *Nautile, Shinkai 6500* and of the ROV *ISIS* during these research cruises. We gratefully acknowledge the chief scientists of the relevant expeditions: Marie-Anne Cambon-Bonavita (Ifremer; BICOSE 2), Ken Takai (JAMSTEC; YK13-05, YK16-E02), Kentaro Nakamura and Satoshi Nakagawa (JAMSTEC; YK09-13) and Jon Copley (NOC; JC82). We also thank Rika Horiuchi (JAMSTEC) for her assistance with μ-CT scanning facilities at JAMSTEC. Faunal collections of *R. exoculata, R. chacei* and *R. kairei* specimens were conducted within international waters. Permission for sampling *R. hybisae* specimens in territorial waters of the Cayman Islands was issued by the appropriate United Kingdom authorities to R/V Yokosuka during YK13-05.

## Data Accessibility

3D Interactive models from each stage and shrimp species are available as supplementary material data. Metadata associated to each shrimp individuals including collection date, sampling location, storage conditions and body size are displayed in Table S1. Surfaces and volumes measurement of shrimp organs obtained from 3D anatomical reconstructions are given in Table S3.

## Author’s contribution

PM collected shrimps during the BICOSE 2 expedition, carried out the identification and measurement of specimens, realized Fluorescent in situ hybridization experiments on *R. hybisae* specimens, conducted data analysis of 3D anatomical reconstruction, lead the conception and design of the study, and drafted the original manuscript; MG conducted FISH analyses on *R. exoculata* and *R. chacei*, and critically revised the manuscript; JTC collected specimens from the JC82 expedition and critically revised the manuscript. HKW collected specimens from YK13-05 and critically revised the manuscript. FP participated in the design and conception of the study and critically revised the manuscript. MACB participated in the design and conception of the study and critically revised the manuscript. CC collected specimens from YK16-06 expeditions, assisted in drafting the original manuscript, participated in the design and conception of the study, and coordinated the study. All authors gave final approval for submission and publication of the manuscript in its current form.

