## supplemental_material.pdf for "Juvenile niches select between two distinct development trajectories and symbiosis modes in vent shrimps"

### Supplementary Figures

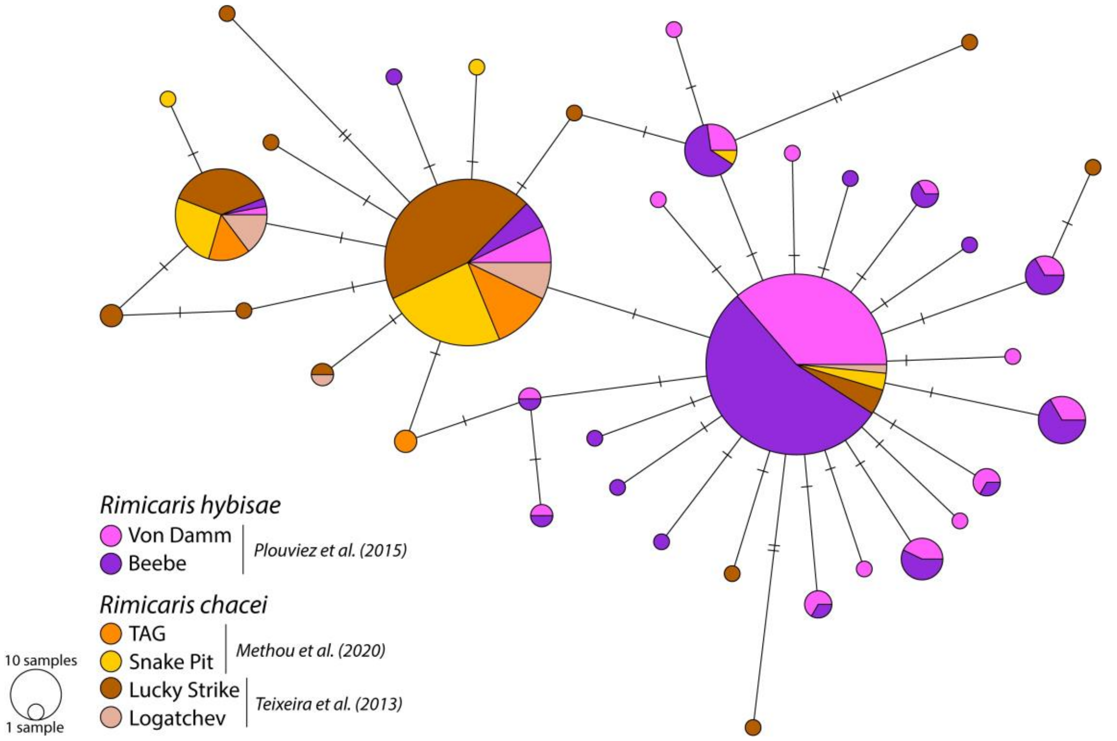


**Figure S1.** COI haplotype network obtained with a median-joining method in PopART (Leigh & Bryant 2015) from published sequences of *R. chacei* (Teixeira et al. 2013, Methou et al. 2020) and *R. hybisae* (Plouviez et al. 2015). Sizes of coloured circles indicate relative haplotype frequencies.

Leigh JW, Bryant D (2015) POPART : full-feature software for haplotype network construction. Methods Ecol Evol 6:1110–1116.

Methou P, Michel LN, Segonzac M, Cambon-Bonavita M-A, Pradillon F (2020) Integrative taxonomy revisits the ontogeny and trophic niches of Rimicaris vent shrimps. R Soc Open Sci 7:200837.

Plouviez S, Jacobson A, Wu M, Van Dover CL (2015) Characterization of vent fauna at the mid-cayman spreading center. Deep Res Part I Oceanogr Res Pap 97:124–133.

Teixeira S, Olu K, Decker C, Cunha RL, Fuchs S, Hourdez S, Serrão EA, Arnaud-Haond S (2013) High connectivity across the fragmented chemosynthetic ecosystems of the deep Atlantic Equatorial Belt: Efficient dispersal mechanisms or questionable endemism? Mol Ecol 22:4663–4680.

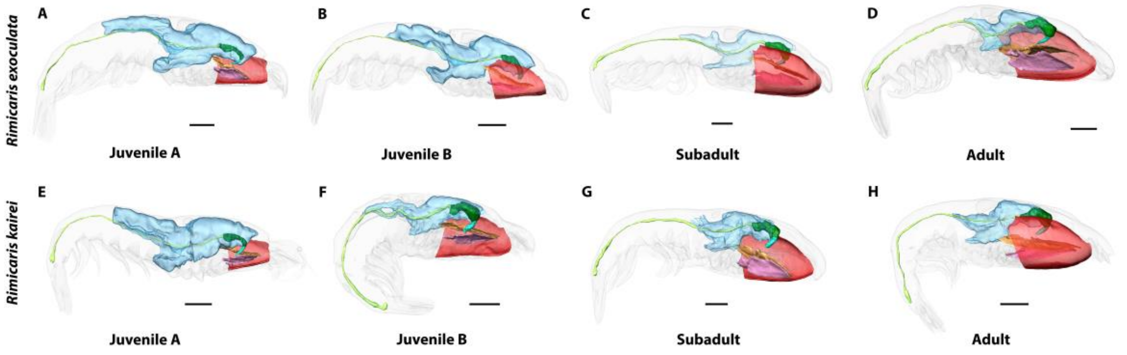


**Figure S2.** 3D anatomical reconstructions of symbiont hosting organs (branchiostegite, scaphognathites, and exopodites) and digestive organs (stomach, hepatopancreas, digestive tube) in *Rimicaris exoculata* (A-D) and *Rimicaris kairei* (E-H) shrimps across post-settlement ontogeny.

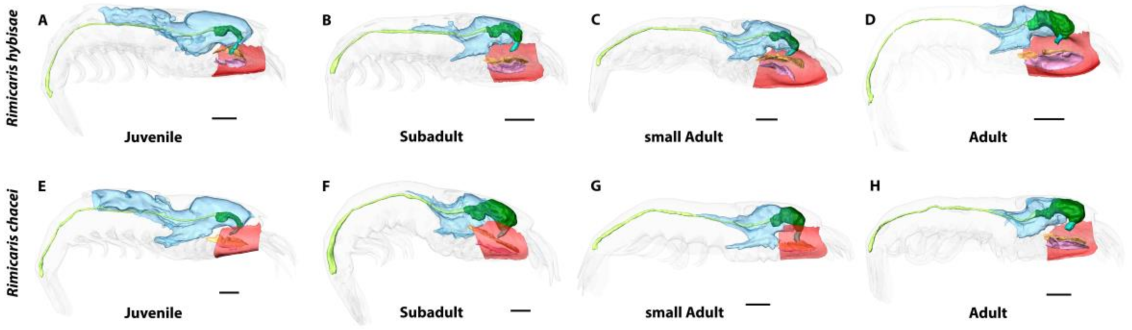


**Figure S3.** 3D anatomical reconstructions of symbiont hosting organs (branchiostegite, scaphognathites, and exopodites) and digestive organs (stomach, hepatopancreas, digestive tube) in *Rimicaris hybisae* (A-D) and *Rimicaris chacei* (E-H) shrimps across post-settlement ontogeny.

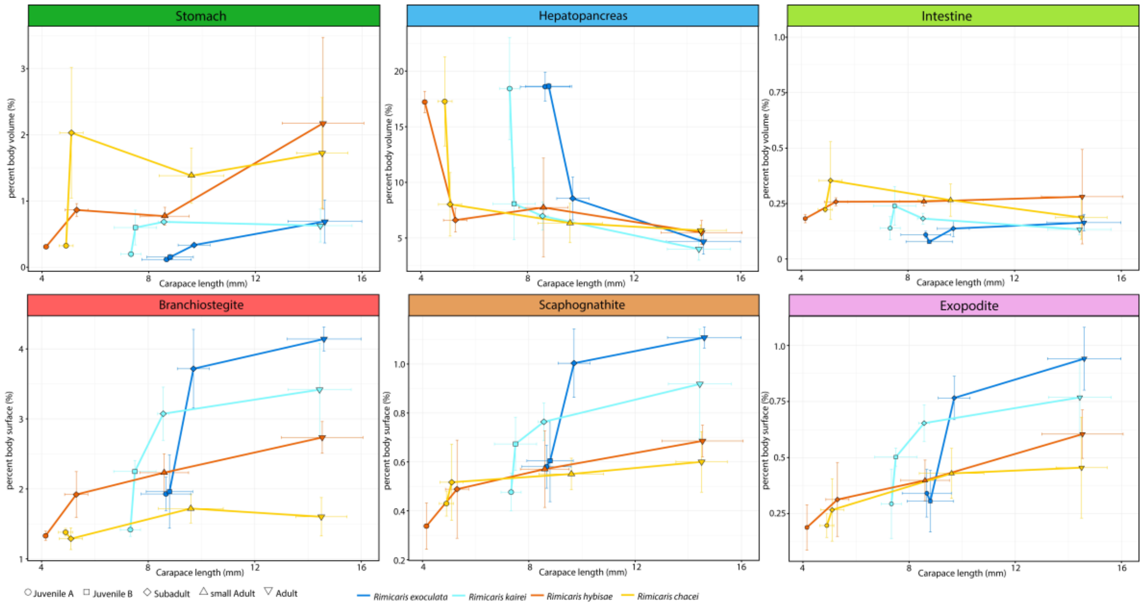


**Figure S4.** Relationship between body size (carapace length) and the relative per cent body volumes or body surfaces of symbiont-hosting organs (branchiostegite, scaphognathites, and exopodites) and digestive organs (stomach, hepatopancreas, digestive tube).
