## supplemental_3Dmodels for "Juvenile niches select between two distinct development trajectories and symbiosis modes in vent shrimps"

### Supplementary Data : Interactive 3D Models

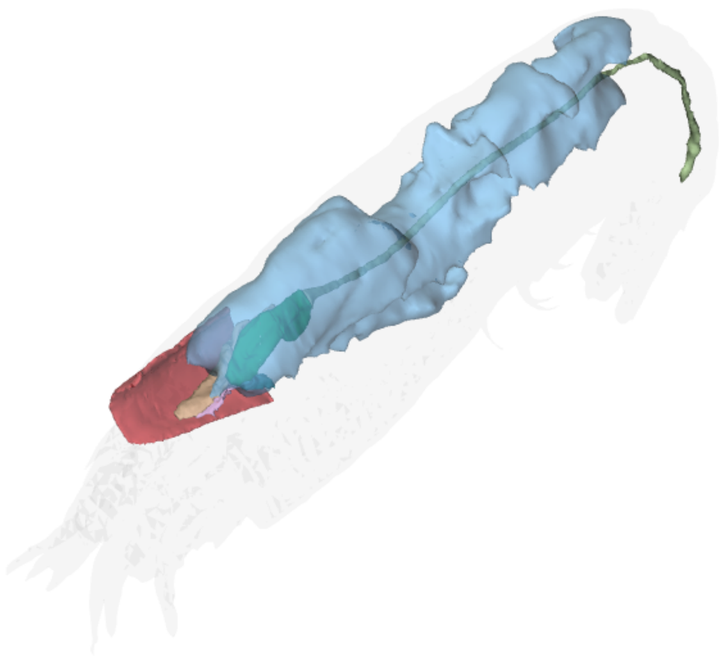

#### *Rimicaris chacei,* Juvenile A

This interactive 3D model can be accessed by clicking on the figure (Adobe Acrobat Reader v7 or higher). Hold left click and drag to rotate, hold down CTRL while doing so to move, and hold down shift while doing do to zoom (alternatively, hold right click and drag or use scroll wheel). Switch between pre-saved views using the drop-down menu in the floating window or click on the 'view' pane in the model tree. Components can also be activated or deactivated by toggling the checkbox in the model tree. The 3D PDF was generated using Adobe Acrobat Pro by importing .u3d files (converted using DAZ Studio) from .obj exports of Amira surface files.

**Supplementary File** for: *Juvenile niches select between two distinct development trajectories and symbiosis modes in vent shrimps* by Pierre Methou, Marion Guéganton, Jonathan T. Copley, Hiromi Kayama Watanabe, Florence Pradillon, Marie-Anne Cambon-Bonavita, and Chong Chen.

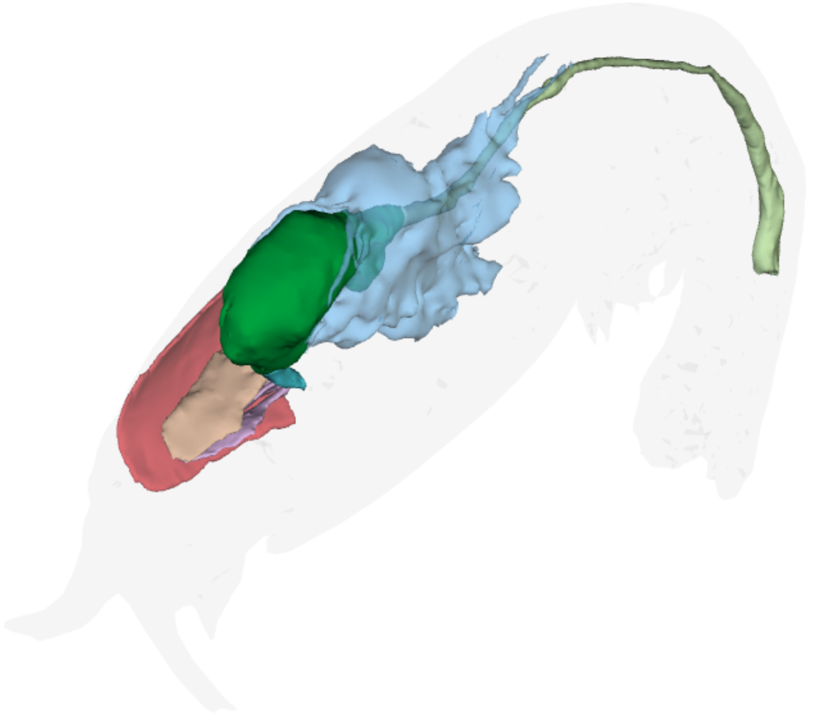

#### *Rimicaris chacei,* Subadult

This interactive 3D model can be accessed by clicking on the figure (Adobe Acrobat Reader v7 or higher). Hold left click and drag to rotate, hold down CTRL while doing so to move, and hold down shift while doing do to zoom (alternatively, hold right click and drag or use scroll wheel). Switch between pre-saved views using the drop-down menu in the floating window or click on the 'view' pane in the model tree. Components can also be activated or deactivated by toggling the checkbox in the model tree. The 3D PDF was generated using Adobe Acrobat Pro by importing .u3d files (converted using DAZ Studio) from .obj exports of Amira surface files.

**Supplementary File** for: *Juvenile niches select between two distinct development trajectories and symbiosis modes in vent shrimps* by Pierre Methou, Marion Guéganton, Jonathan T. Copley, Hiromi Kayama Watanabe, Florence Pradillon, Marie-Anne Cambon-Bonavita, and Chong Chen.

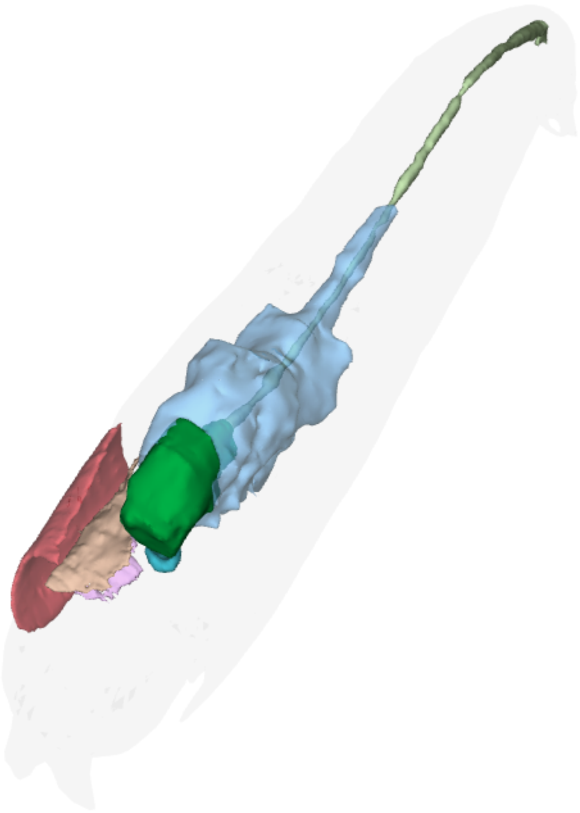

#### *Rimicaris chacei*, small Adult

This interactive 3D model can be accessed by clicking on the figure (Adobe Acrobat Reader v7 or higher). Hold left click and drag to rotate, hold down CTRL while doing so to move, and hold down shift while doing do to zoom (alternatively, hold right click and drag or use scroll wheel). Switch between pre-saved views using the drop-down menu in the floating window or click on the 'view' pane in the model tree. Components can also be activated or deactivated by toggling the checkbox in the model tree. The 3D PDF was generated using Adobe Acrobat Pro by importing .u3d files (converted using DAZ Studio) from .obj exports of Amira surface files.

**Supplementary File** for: *Juvenile niches select between two distinct development trajectories and symbiosis modes in vent shrimps* by Pierre Methou, Marion Guéganton, Jonathan T. Copley, Hiromi Kayama Watanabe, Florence Pradillon, Marie-Anne Cambon-Bonavita, and Chong Chen.

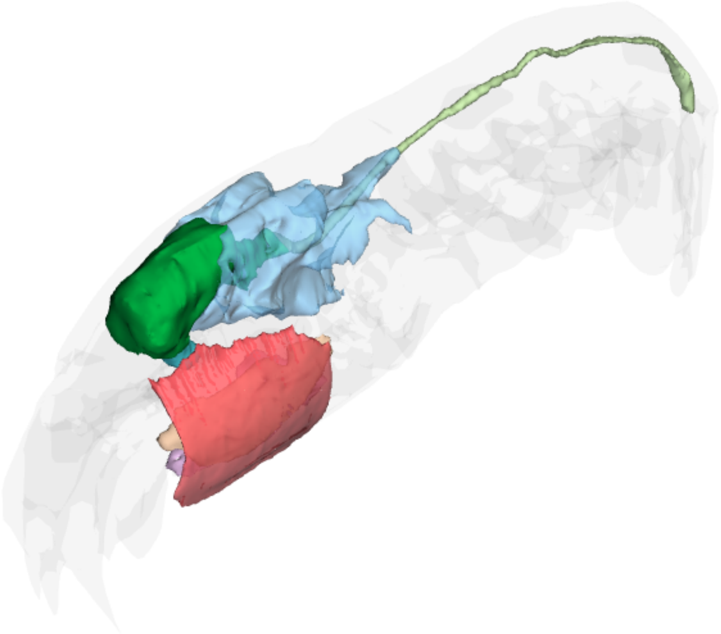

#### *Rimicaris chacei*, Adult

This interactive 3D model can be accessed by clicking on the figure (Adobe Acrobat Reader v7 or higher). Hold left click and drag to rotate, hold down CTRL while doing so to move, and hold down shift while doing do to zoom (alternatively, hold right click and drag or use scroll wheel). Switch between pre-saved views using the drop-down menu in the floating window or click on the 'view' pane in the model tree. Components can also be activated or deactivated by toggling the checkbox in the model tree. The 3D PDF was generated using Adobe Acrobat Pro by importing .u3d files (converted using DAZ Studio) from .obj exports of Amira surface files.

**Supplementary File** for: *Juvenile niches select between two distinct development trajectories and symbiosis modes in vent shrimps* by Pierre Methou, Marion Guéganton, Jonathan T. Copley, Hiromi Kayama Watanabe, Florence Pradillon, Marie-Anne Cambon-Bonavita, and Chong Chen.

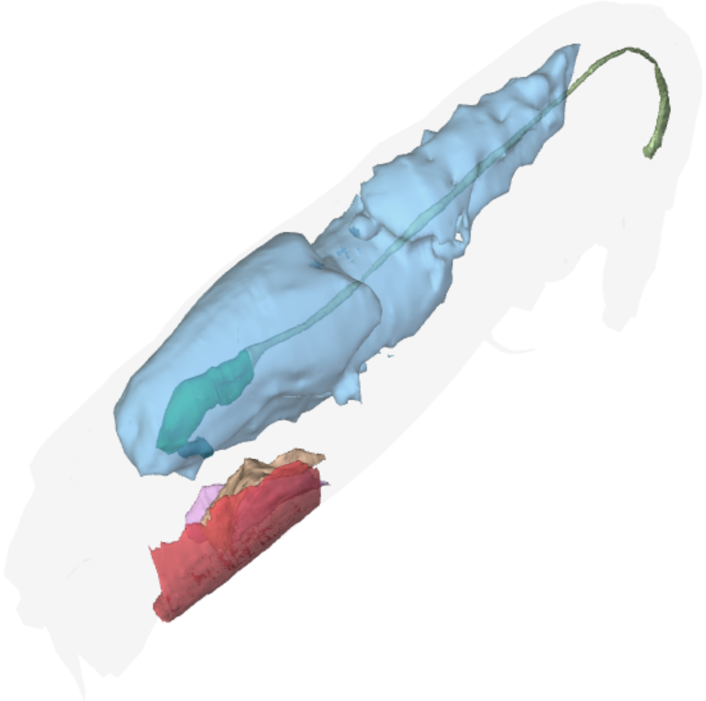

#### *Rimicaris hybisae*, Juvenile A

This interactive 3D model can be accessed by clicking on the figure (Adobe Acrobat Reader v7 or higher). Hold left click and drag to rotate, hold down CTRL while doing so to move, and hold down shift while doing do to zoom (alternatively, hold right click and drag or use scroll wheel). Switch between pre-saved views using the drop-down menu in the floating window or click on the 'view' pane in the model tree. Components can also be activated or deactivated by toggling the checkbox in the model tree. The 3D PDF was generated using Adobe Acrobat Pro by importing .u3d files (converted using DAZ Studio) from .obj exports of Amira surface files.

**Supplementary File** for: *Juvenile niches select between two distinct development trajectories and symbiosis modes in vent shrimps* by Pierre Methou, Marion Guéganton, Jonathan T. Copley, Hiromi Kayama Watanabe, Florence Pradillon, Marie-Anne Cambon-Bonavita, and Chong Chen.

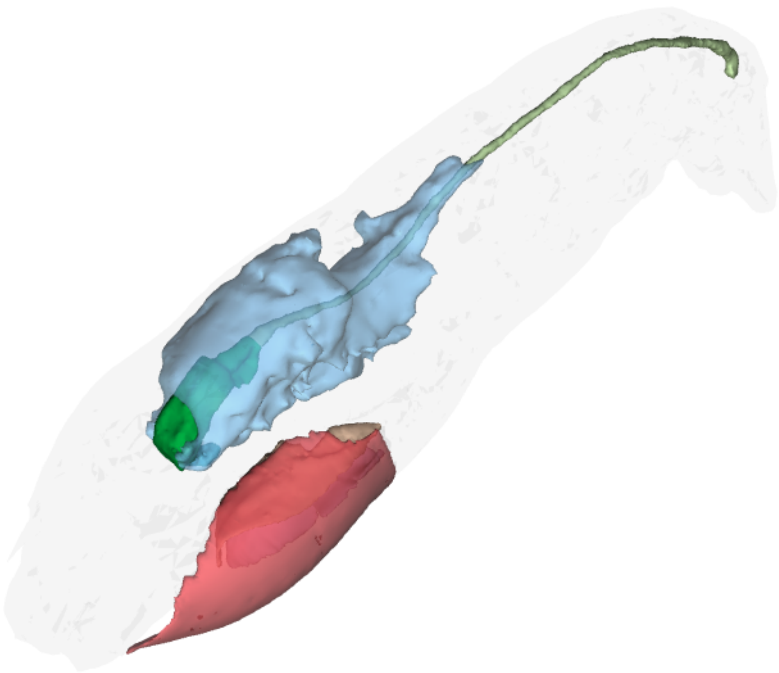

#### *Rimicaris hybisae,* small Adult

This interactive 3D model can be accessed by clicking on the figure (Adobe Acrobat Reader v7 or higher). Hold left click and drag to rotate, hold down CTRL while doing so to move, and hold down shift while doing do to zoom (alternatively, hold right click and drag or use scroll wheel). Switch between pre-saved views using the drop-down menu in the floating window or click on the 'view' pane in the model tree. Components can also be activated or deactivated by toggling the checkbox in the model tree. The 3D PDF was generated using Adobe Acrobat Pro by importing .u3d files (converted using DAZ Studio) from .obj exports of Amira surface files.

**Supplementary File** for: *Juvenile niches select between two distinct development trajectories and symbiosis modes in vent shrimps* by Pierre Methou, Marion Guéganton, Jonathan T. Copley, Hiromi Kayama Watanabe, Florence Pradillon, Marie-Anne Cambon-Bonavita, and Chong Chen.

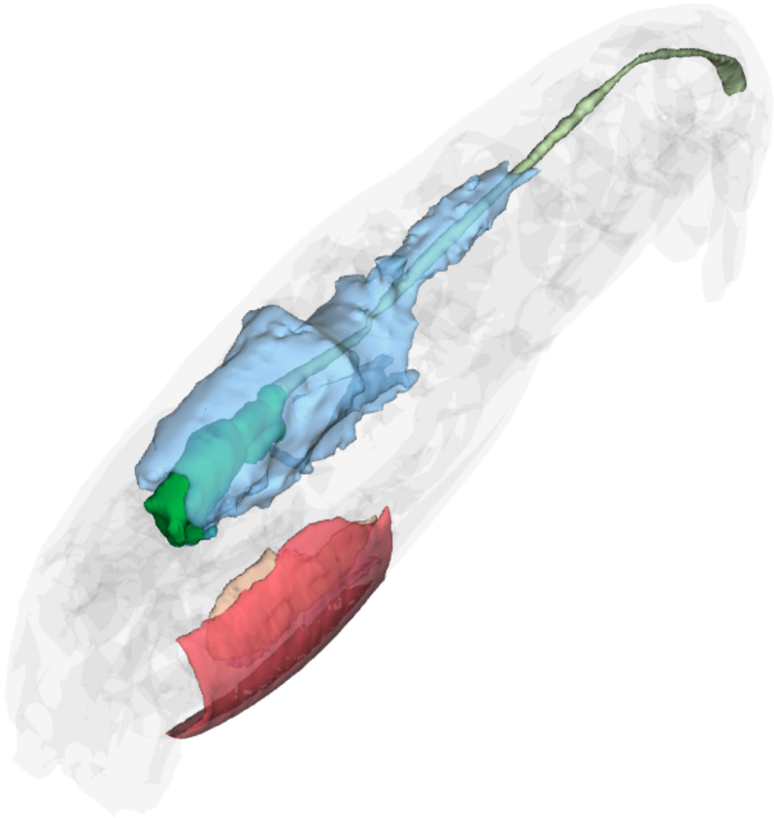

#### *Rimicaris hybisae*, Subadult

This interactive 3D model can be accessed by clicking on the figure (Adobe Acrobat Reader v7 or higher). Hold left click and drag to rotate, hold down CTRL while doing so to move, and hold down shift while doing do to zoom (alternatively, hold right click and drag or use scroll wheel). Switch between pre-saved views using the drop-down menu in the floating window or click on the 'view' pane in the model tree. Components can also be activated or deactivated by toggling the checkbox in the model tree. The 3D PDF was generated using Adobe Acrobat Pro by importing .u3d files (converted using DAZ Studio) from .obj exports of Amira surface files.

**Supplementary File** for: *Juvenile niches select between two distinct development trajectories and symbiosis modes in vent shrimps* by Pierre Methou, Marion Guéganton, Jonathan T. Copley, Hiromi Kayama Watanabe, Florence Pradillon, Marie-Anne Cambon-Bonavita, and Chong Chen

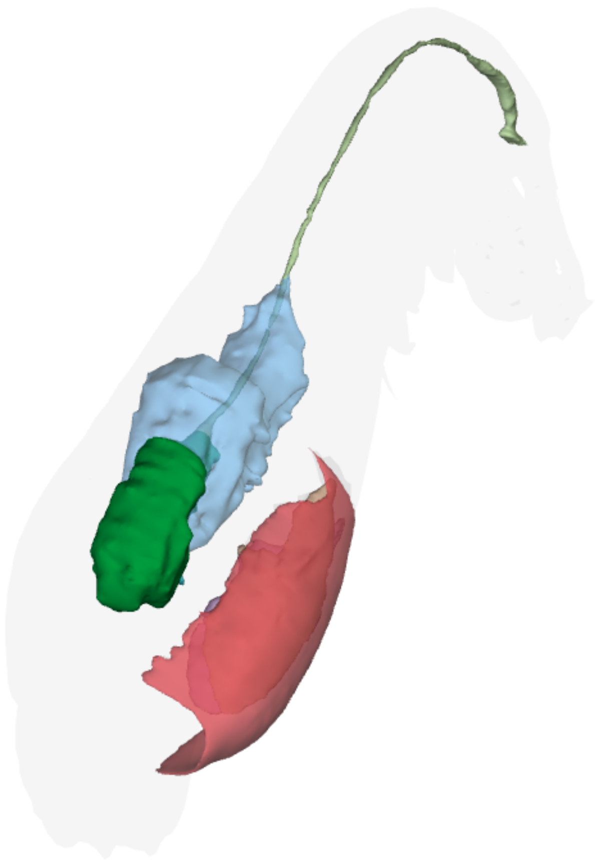

#### *Rimicaris hybisae*, Adult

This interactive 3D model can be accessed by clicking on the figure (Adobe Acrobat Reader v7 or higher). Hold left click and drag to rotate, hold down CTRL while doing so to move, and hold down shift while doing do to zoom (alternatively, hold right click and drag or use scroll wheel). Switch between pre-saved views using the drop-down menu in the floating window or click on the 'view' pane in the model tree. Components can also be activated or deactivated by toggling the checkbox in the model tree. The 3D PDF was generated using Adobe Acrobat Pro by importing .u3d files (converted using DAZ Studio) from .obj exports of Amira surface files.

**Supplementary File** for: *Juvenile niches select between two distinct development trajectories and symbiosis modes in vent shrimps* by Pierre Methou, Marion Guéganton, Jonathan T. Copley, Hiromi Kayama Watanabe, Florence Pradillon, Marie-Anne Cambon-Bonavita, and Chong Chen.

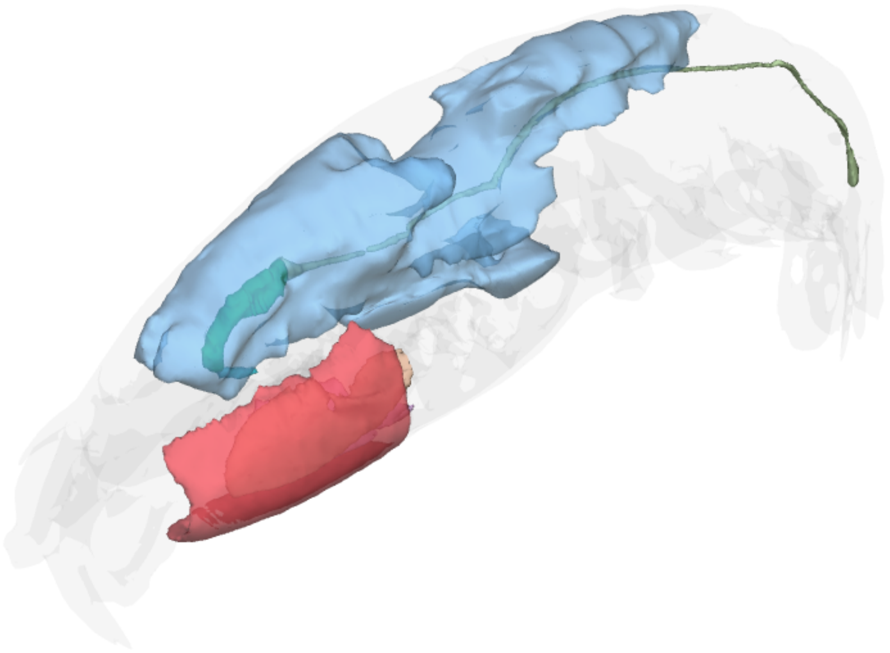

#### *Rimicaris exoculata*, Juvenile A

This interactive 3D model can be accessed by clicking on the figure (Adobe Acrobat Reader v7 or higher). Hold left click and drag to rotate, hold down CTRL while doing so to move, and hold down shift while doing do to zoom (alternatively, hold right click and drag or use scroll wheel). Switch between pre-saved views using the drop-down menu in the floating window or click on the 'view' pane in the model tree. Components can also be activated or deactivated by toggling the checkbox in the model tree. The 3D PDF was generated using Adobe Acrobat Pro by importing .u3d files (converted using DAZ Studio) from .obj exports of Amira surface files.

**Supplementary File** for: *Juvenile niches select between two distinct development trajectories and symbiosis modes in vent shrimps* by Pierre Methou, Marion Guéganton, Jonathan T. Copley, Hiromi Kayama Watanabe, Florence Pradillon, Marie-Anne Cambon-Bonavita, and Chong Chen.

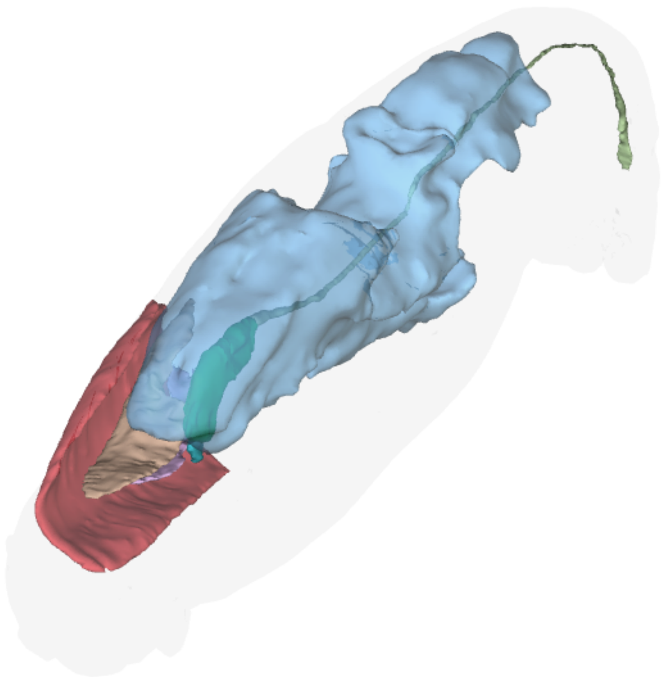

#### *Rimicaris exoculata*, Juvenile B

This interactive 3D model can be accessed by clicking on the figure (Adobe Acrobat Reader v7 or higher). Hold left click and drag to rotate, hold down CTRL while doing so to move, and hold down shift while doing do to zoom (alternatively, hold right click and drag or use scroll wheel). Switch between pre-saved views using the drop-down menu in the floating window or click on the 'view' pane in the model tree. Components can also be activated or deactivated by toggling the checkbox in the model tree. The 3D PDF was generated using Adobe Acrobat Pro by importing .u3d files (converted using DAZ Studio) from .obj exports of Amira surface files.

**Supplementary File** for: *Juvenile niches select between two distinct development trajectories and symbiosis modes in vent shrimps* by Pierre Methou, Marion Guéganton, Jonathan T. Copley, Hiromi Kayama Watanabe, Florence Pradillon, Marie-Anne Cambon-Bonavita, and Chong Chen.

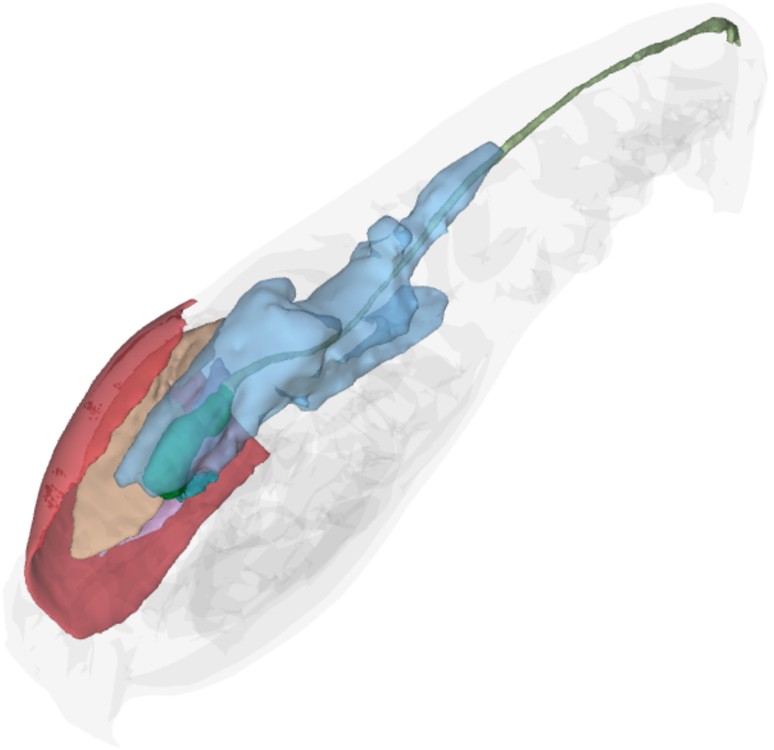

#### *Rimicaris exoculata*, Subadult

This interactive 3D model can be accessed by clicking on the figure (Adobe Acrobat Reader v7 or higher). Hold left click and drag to rotate, hold down CTRL while doing so to move, and hold down shift while doing do to zoom (alternatively, hold right click and drag or use scroll wheel). Switch between pre-saved views using the drop-down menu in the floating window or click on the 'view' pane in the model tree. Components can also be activated or deactivated by toggling the checkbox in the model tree. The 3D PDF was generated using Adobe Acrobat Pro by importing .u3d files (converted using DAZ Studio) from .obj exports of Amira surface files.

**Supplementary File** for: *Juvenile niches select between two distinct development trajectories and symbiosis modes in vent shrimps* by Pierre Methou, Marion Guéganton, Jonathan T. Copley, Hiromi Kayama Watanabe, Florence Pradillon, Marie-Anne Cambon-Bonavita, and Chong Chen

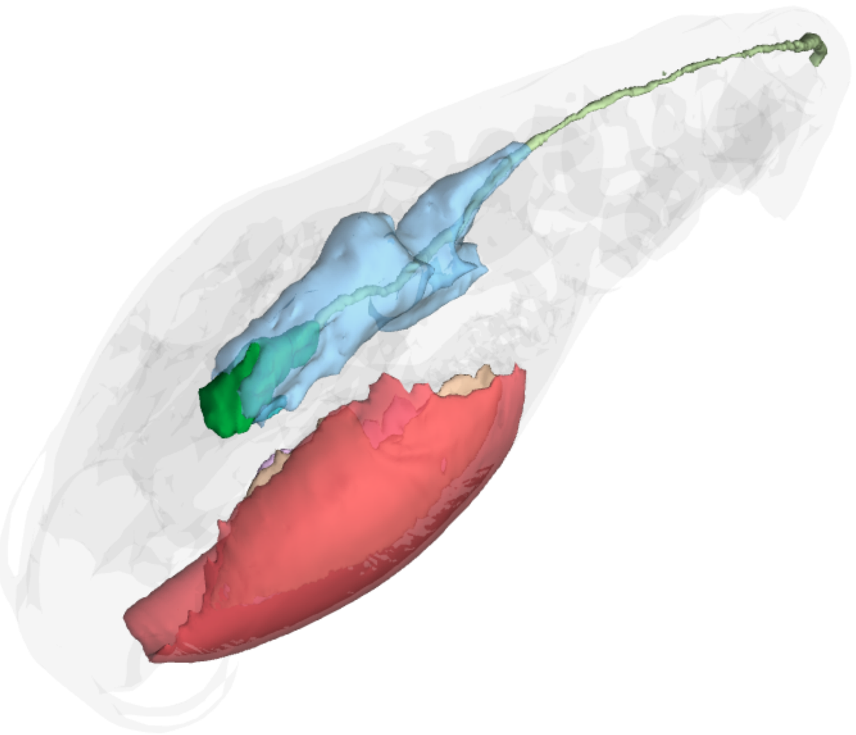

#### *Rimicaris exoculata,* Adult

This interactive 3D model can be accessed by clicking on the figure (Adobe Acrobat Reader v7 or higher). Hold left click and drag to rotate, hold down CTRL while doing so to move, and hold down shift while doing do to zoom (alternatively, hold right click and drag or use scroll wheel). Switch between pre-saved views using the drop-down menu in the floating window or click on the 'view' pane in the model tree. Components can also be activated or deactivated by toggling the checkbox in the model tree. The 3D PDF was generated using Adobe Acrobat Pro by importing .u3d files (converted using DAZ Studio) from .obj exports of Amira surface files.

**Supplementary File** for: *Juvenile niches select between two distinct development trajectories and symbiosis modes in vent shrimps* by Pierre Methou, Marion Guéganton, Jonathan T. Copley, Hiromi Kayama Watanabe, Florence Pradillon, Marie-Anne Cambon-Bonavita, and Chong Chen.

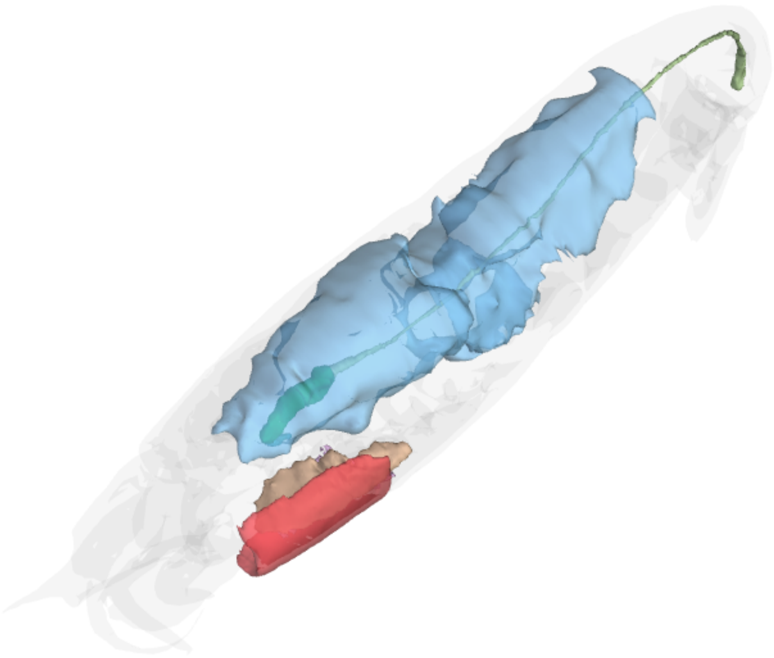

#### *Rimicaris kairei*, Juvenile A

This interactive 3D model can be accessed by clicking on the figure (Adobe Acrobat Reader v7 or higher). Hold left click and drag to rotate, hold down CTRL while doing so to move, and hold down shift while doing do to zoom (alternatively, hold right click and drag or use scroll wheel). Switch between pre-saved views using the drop-down menu in the floating window or click on the 'view' pane in the model tree. Components can also be activated or deactivated by toggling the checkbox in the model tree. The 3D PDF was generated using Adobe Acrobat Pro by importing .u3d files (converted using DAZ Studio) from .obj exports of Amira surface files.

**Supplementary File** for: *Juvenile niches select between two distinct development trajectories and symbiosis modes in vent shrimps* by Pierre Methou, Marion Guéganton, Jonathan T. Copley, Hiromi Kayama Watanabe, Florence Pradillon, Marie-Anne Cambon-Bonavita, and Chong Chen.

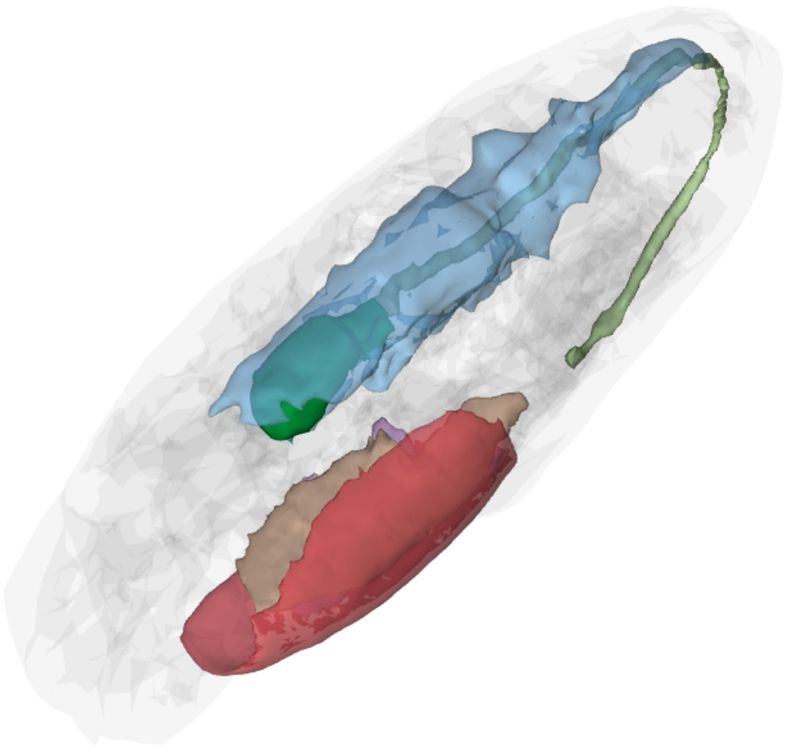

#### *Rimicaris kairei*, Juvenile B

This interactive 3D model can be accessed by clicking on the figure (Adobe Acrobat Reader v7 or higher). Hold left click and drag to rotate, hold down CTRL while doing so to move, and hold down shift while doing do to zoom (alternatively, hold right click and drag or use scroll wheel). Switch between pre-saved views using the drop-down menu in the floating window or click on the 'view' pane in the model tree. Components can also be activated or deactivated by toggling the checkbox in the model tree. The 3D PDF was generated using Adobe Acrobat Pro by importing .u3d files (converted using DAZ Studio) from .obj exports of Amira surface files.

**Supplementary File** for: *Juvenile niches select between two distinct development trajectories and symbiosis modes in vent shrimps* by Pierre Methou, Marion Guéganton, Jonathan T. Copley, Hiromi Kayama Watanabe, Florence Pradillon, Marie-Anne Cambon-Bonavita, and Chong Chen.

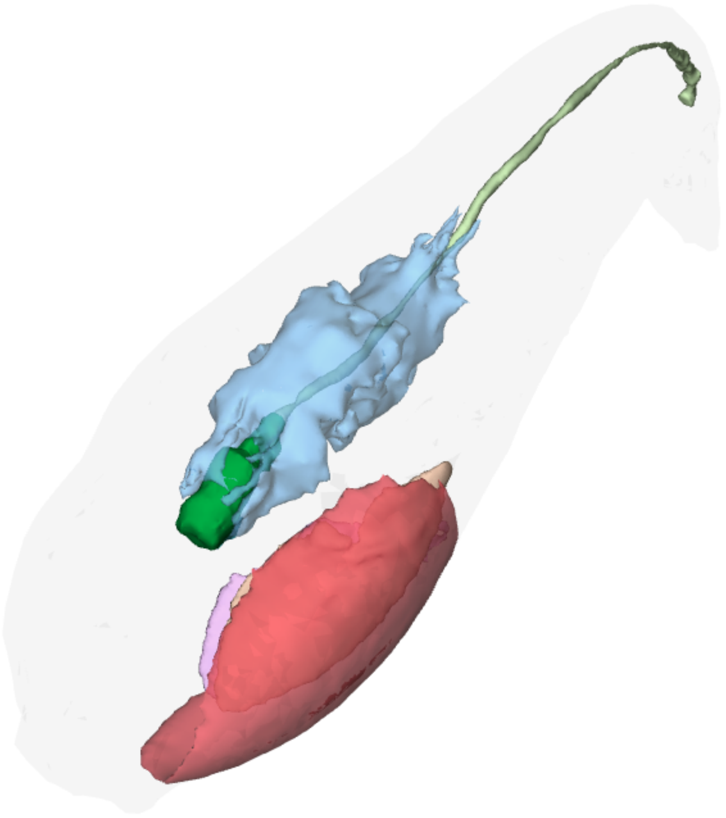

#### *Rimicaris kairei,* Subadult

This interactive 3D model can be accessed by clicking on the figure (Adobe Acrobat Reader v7 or higher). Hold left click and drag to rotate, hold down CTRL while doing so to move, and hold down shift while doing do to zoom (alternatively, hold right click and drag or use scroll wheel). Switch between pre-saved views using the drop-down menu in the floating window or click on the 'view' pane in the model tree. Components can also be activated or deactivated by toggling the checkbox in the model tree. The 3D PDF was generated using Adobe Acrobat Pro by importing .u3d files (converted using DAZ Studio) from .obj exports of Amira surface files.

**Supplementary File** for: *Juvenile niches select between two distinct development trajectories and symbiosis modes in vent shrimps* by Pierre Methou, Marion Guéganton, Jonathan T. Copley, Hiromi Kayama Watanabe, Florence Pradillon, Marie-Anne Cambon-Bonavita, and Chong Chen.

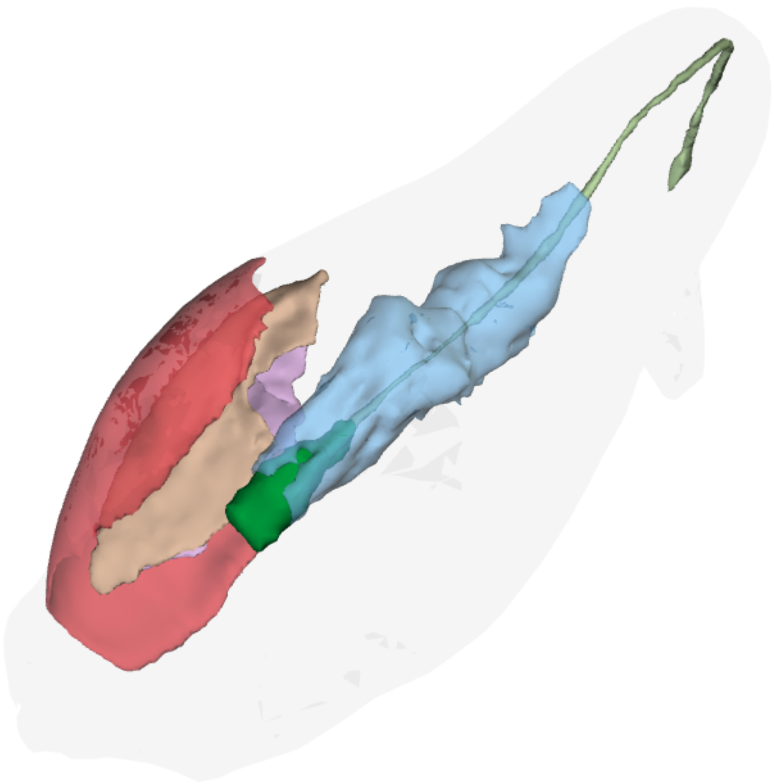

#### *Rimicaris kairei,* Adult

This interactive  3D model can be accessed by clicking on the figure (Adobe Acrobat Reader v7 or higher). Hold left click and drag to rotate, hold down CTRL while doing so to move, and hold down shift while doing do to zoom (alternatively, hold right click and drag or use scroll wheel). Switch between pre-saved views using the drop-down menu in the floating window or click on the 'view' pane in the model tree. Components can also be activated or deactivated by toggling the checkbox in the model tree. The 3D PDF was generated using Adobe Acrobat Pro by importing .u3d files (converted using DAZ Studio) from .obj exports of Amira surface files.

**Supplementary File** for: *Juvenile niches select between two distinct development trajectories and symbiosis modes in vent shrimps* by Pierre Methou, Marion Guéganton, Jonathan T. Copley, Hiromi Kayama Watanabe, Florence Pradillon, Marie-Anne Cambon-Bonavita, and Chong Chen.
